## Supplementary figures and images for "Phylogenetic diversity analysis of shotgun metagenomic reads describes gut microbiome development and treatment effects in the post-weaned pig"

### Supplementary Figure 1

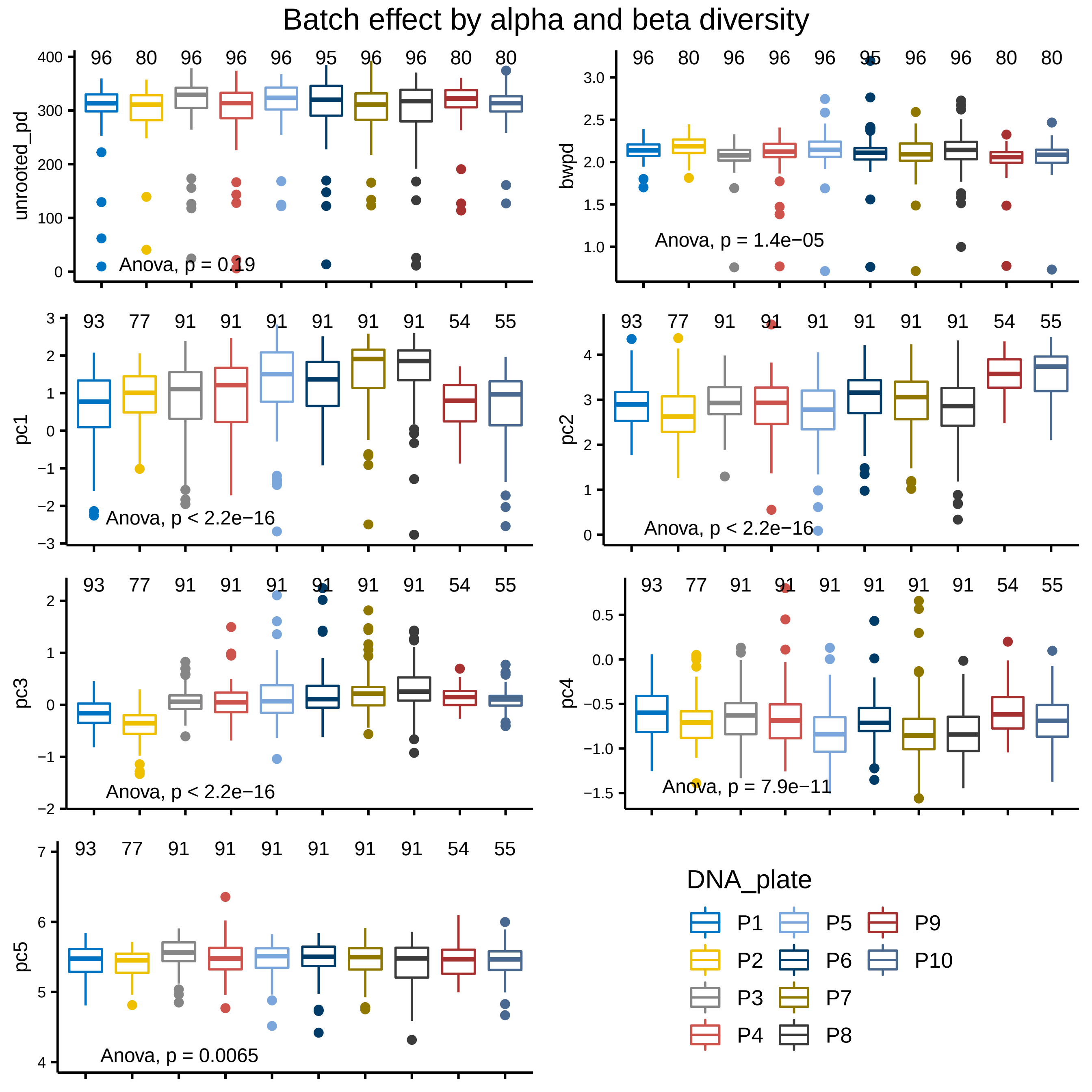

### Supplementary Figure 3

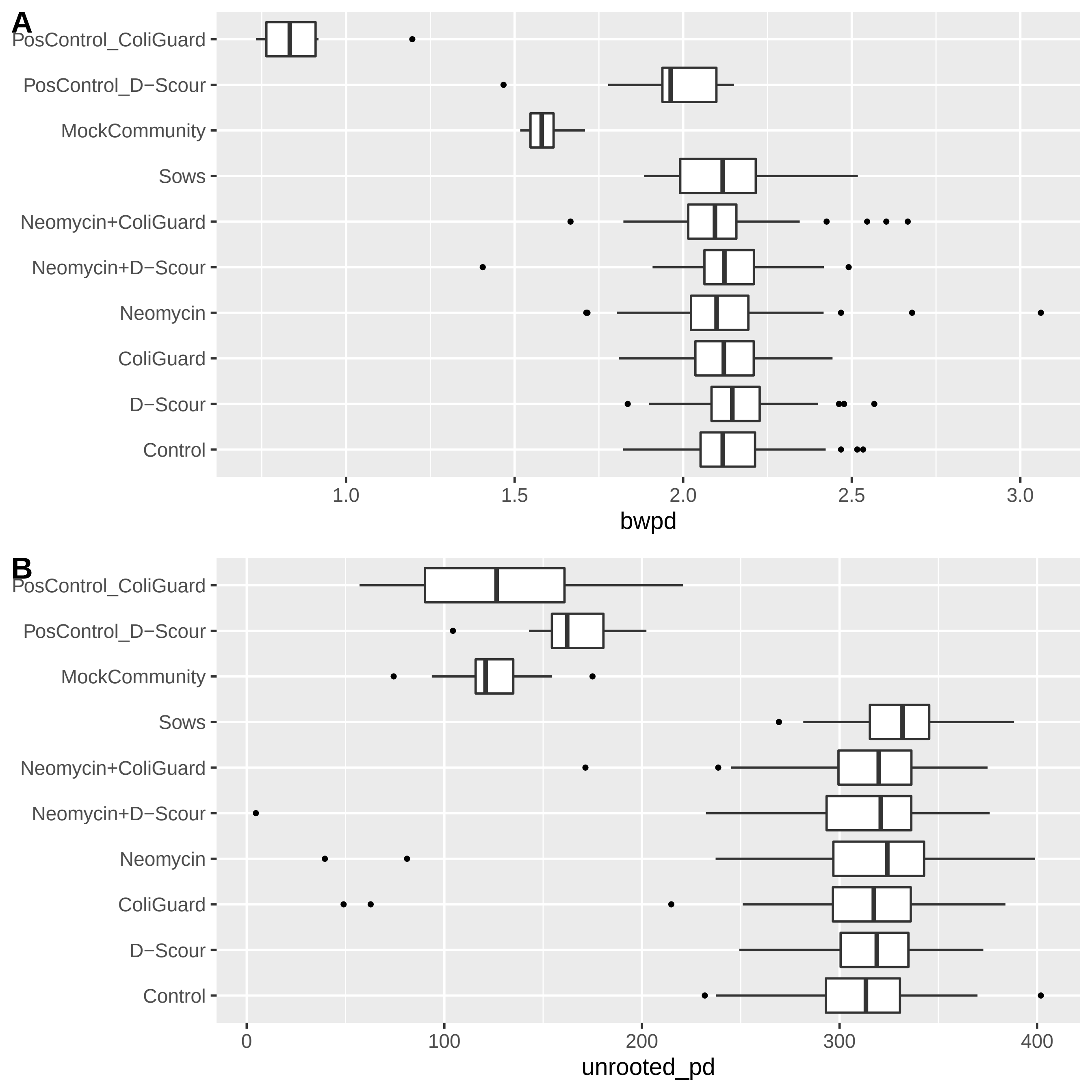

### Supplementary Figure 3

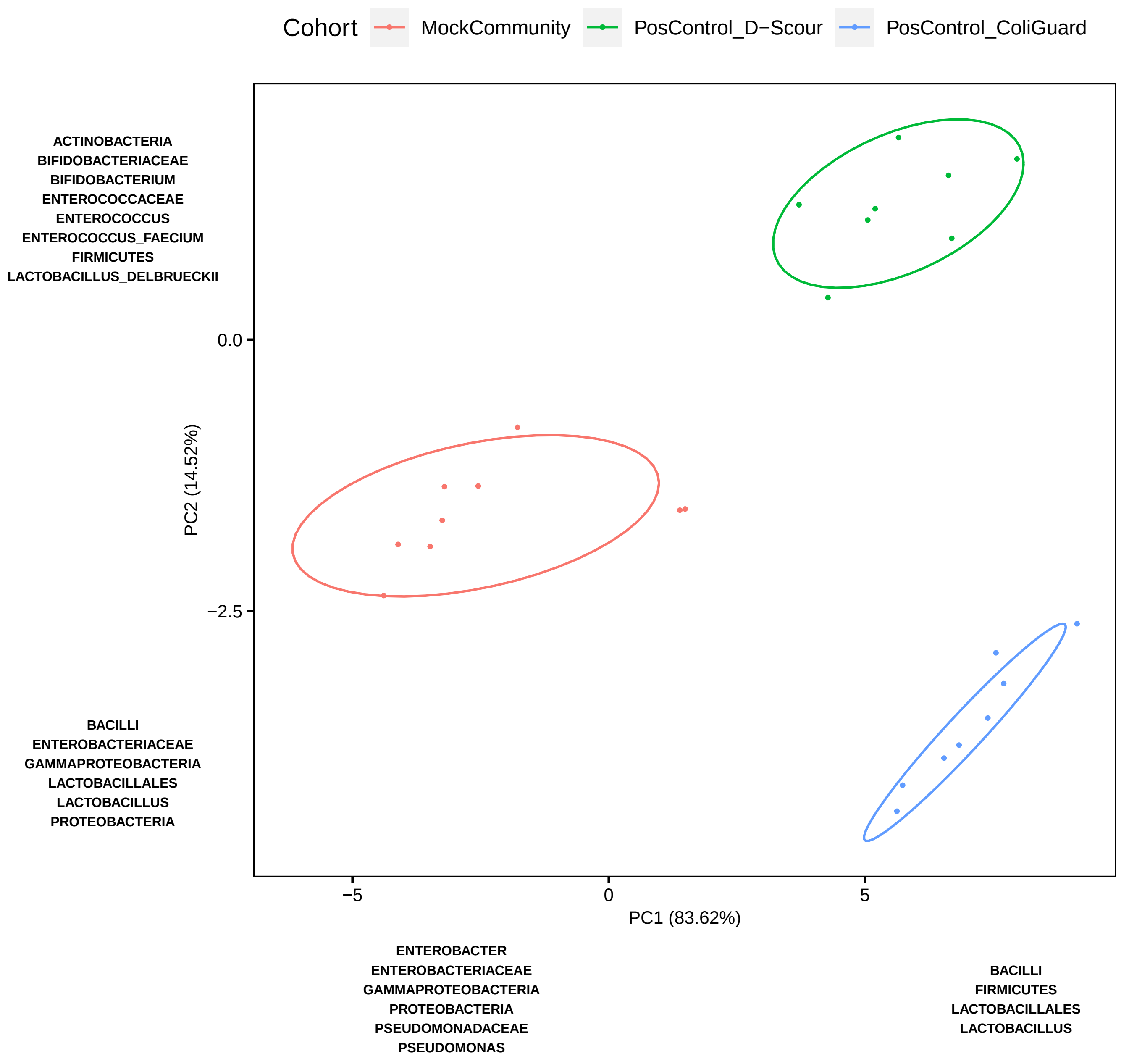

### Supplementary Figure 4

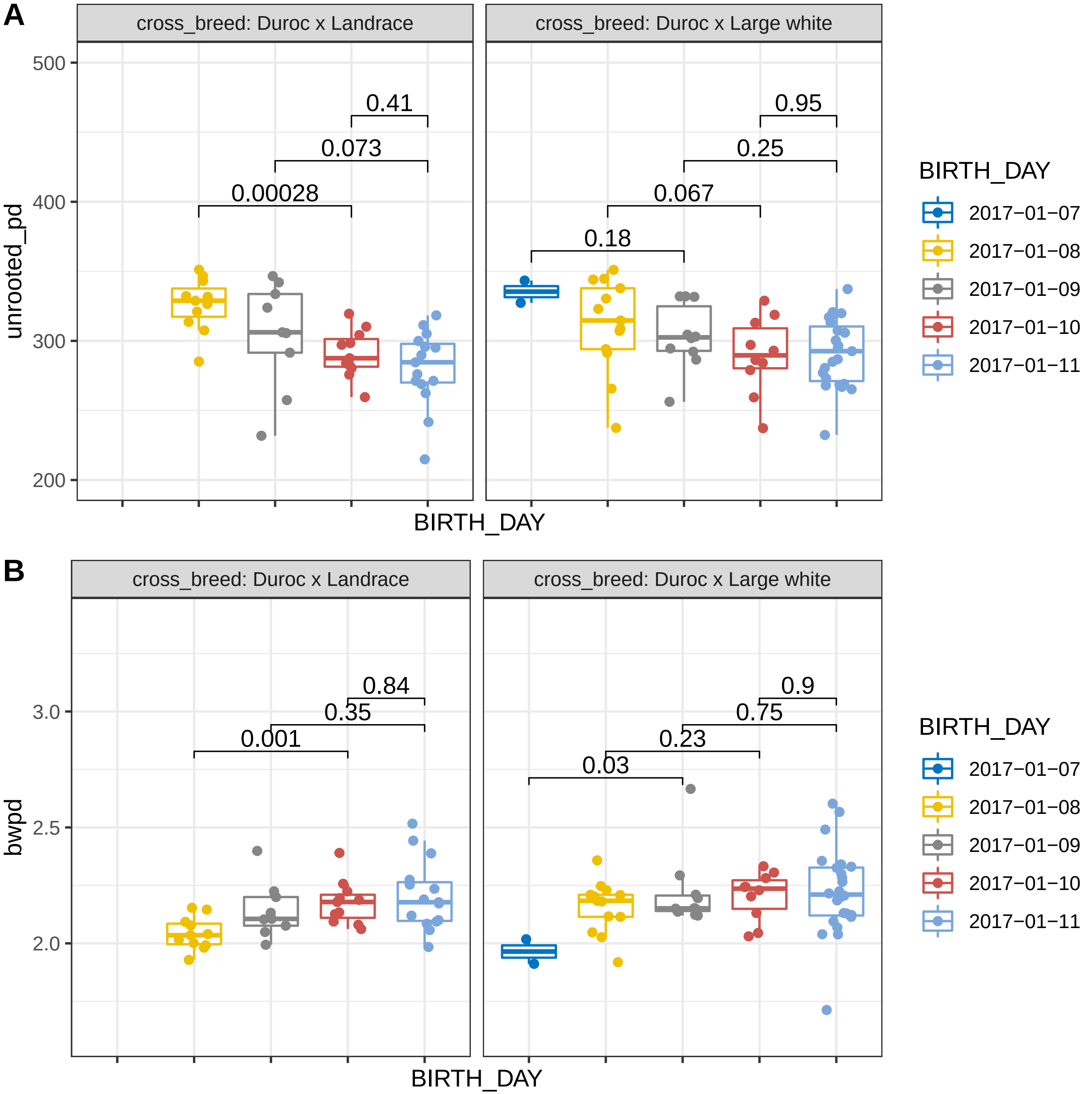

### Supplementary Figure 5

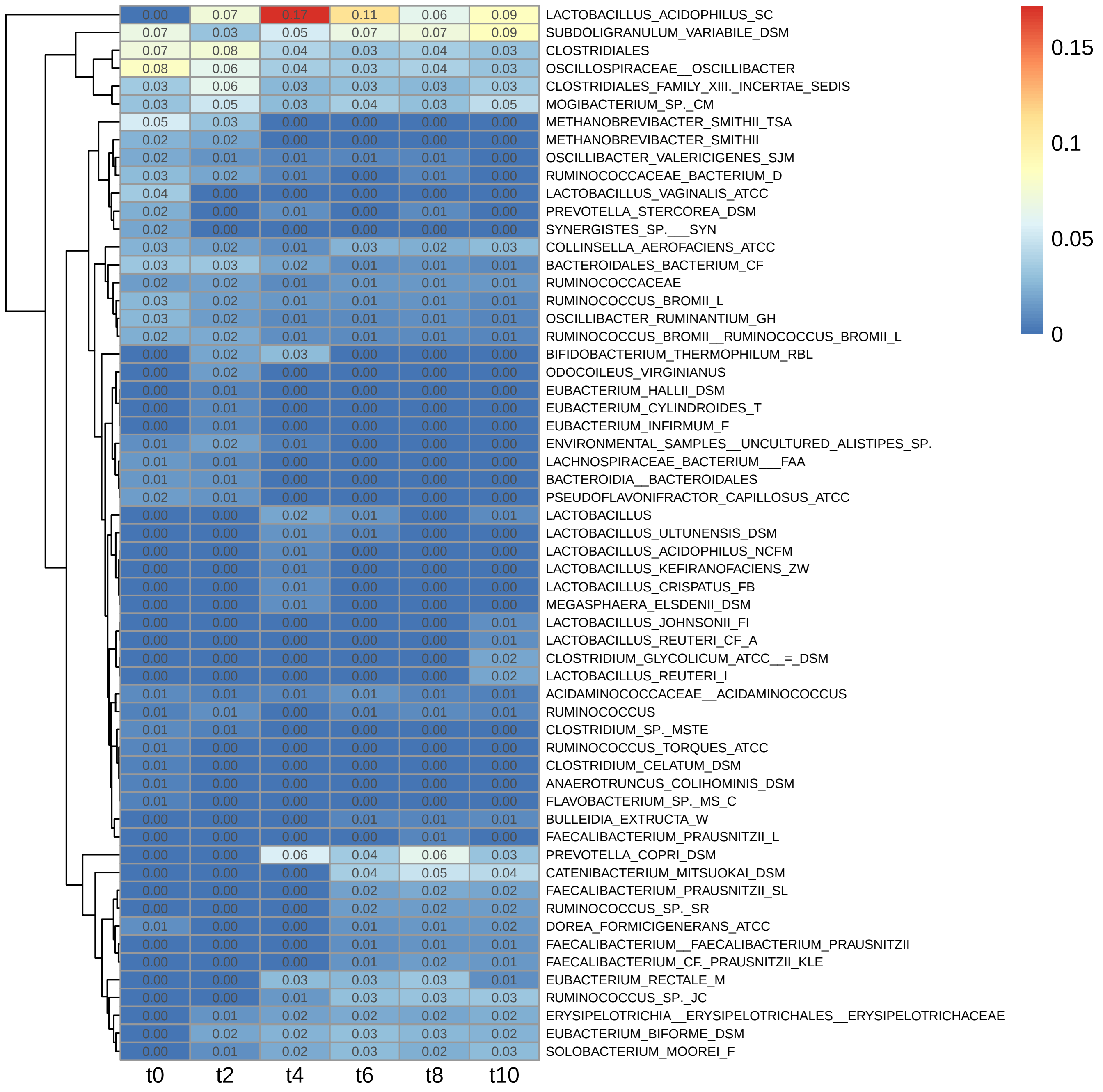

### Supplementary Figure 6

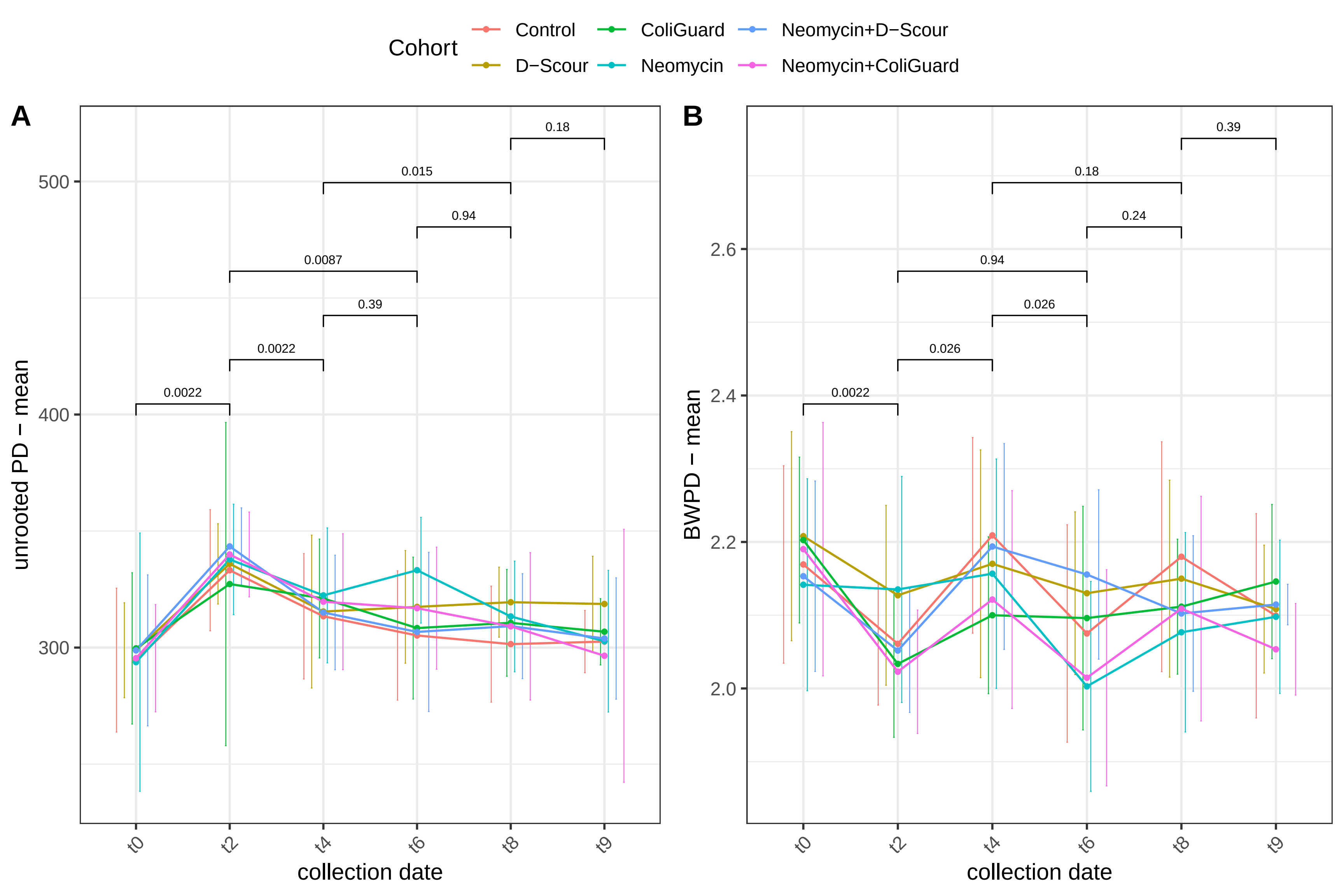

### Supplementary Figure 7

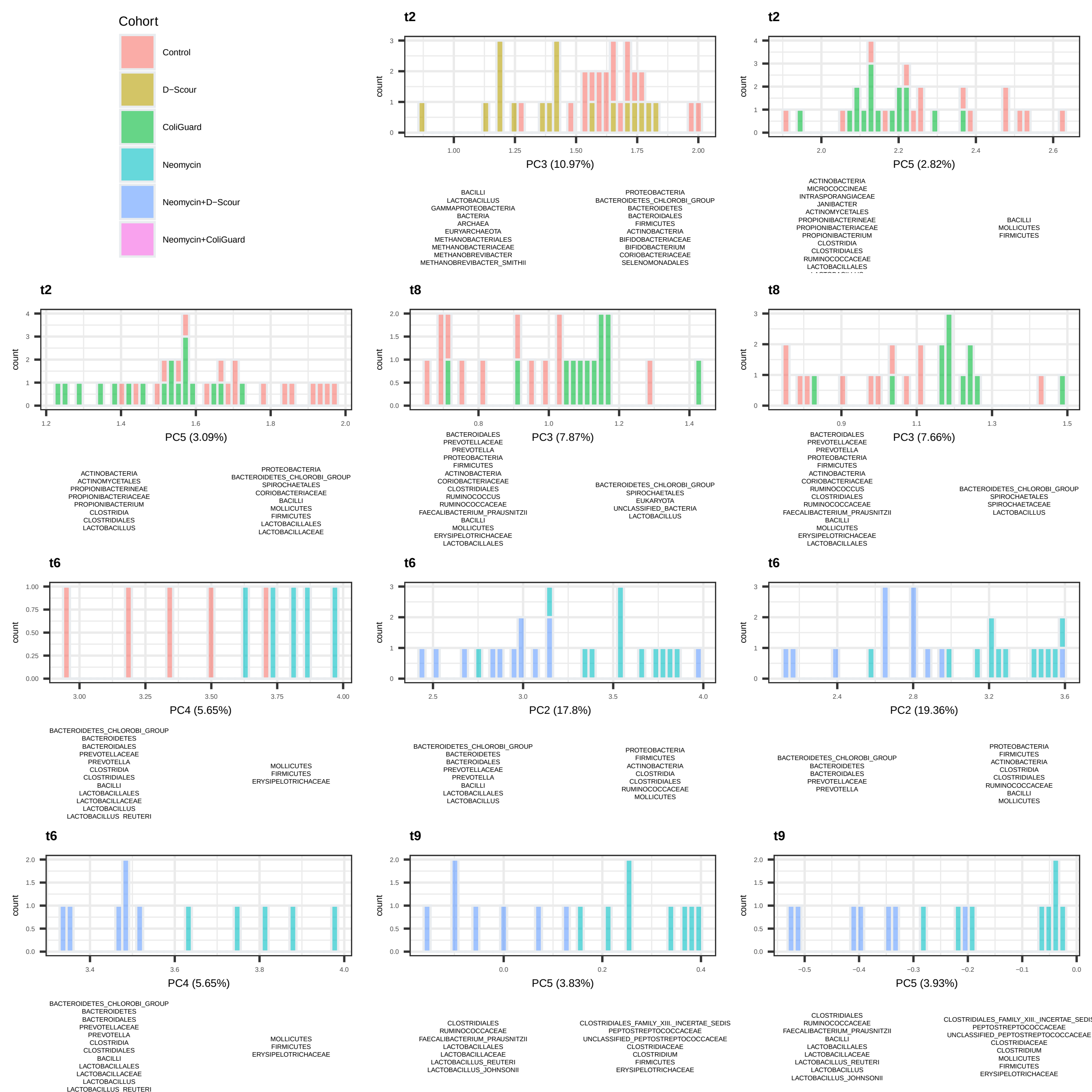

### Supplementary Figure 8

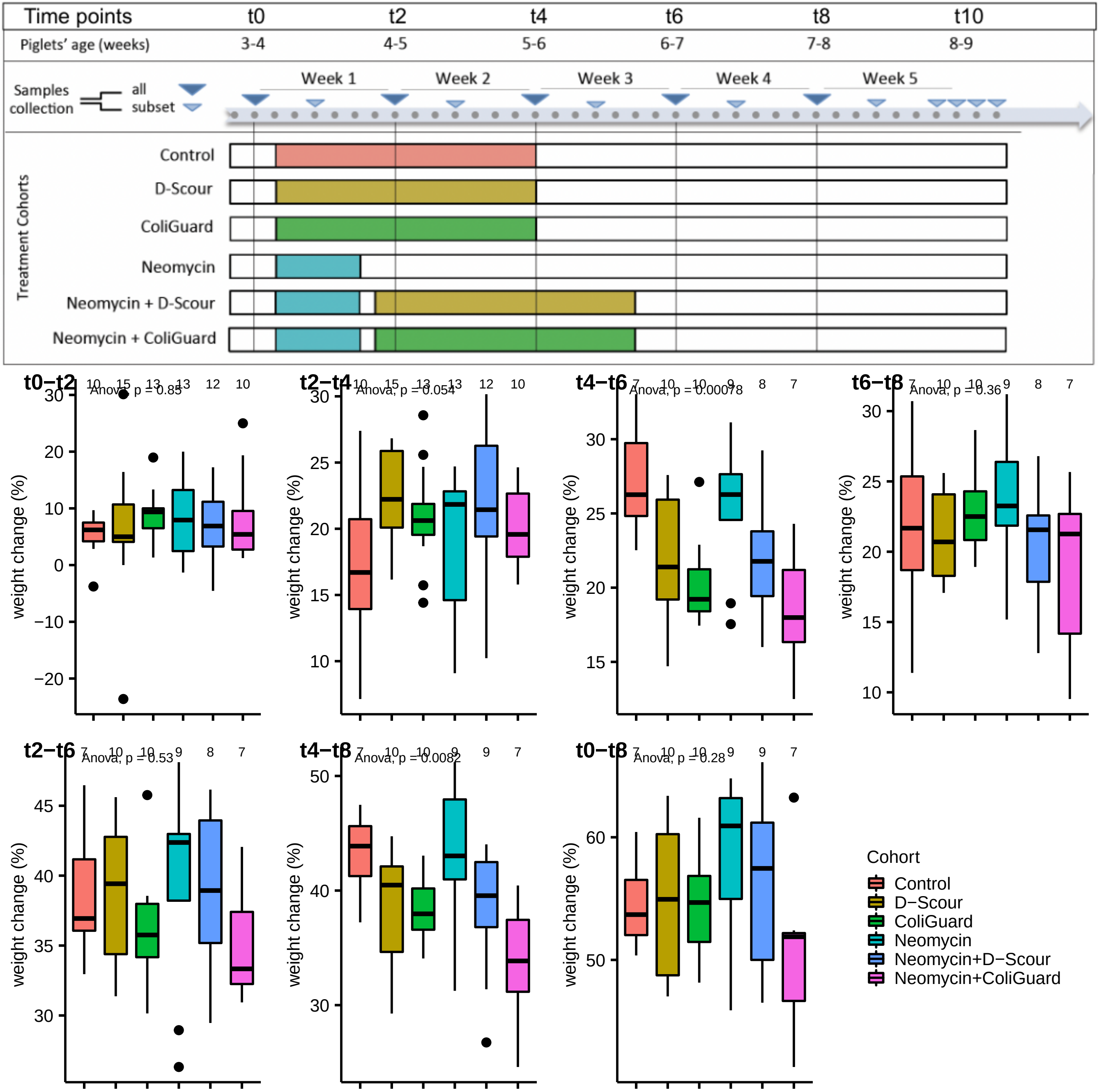

### Supplementary Figure 9

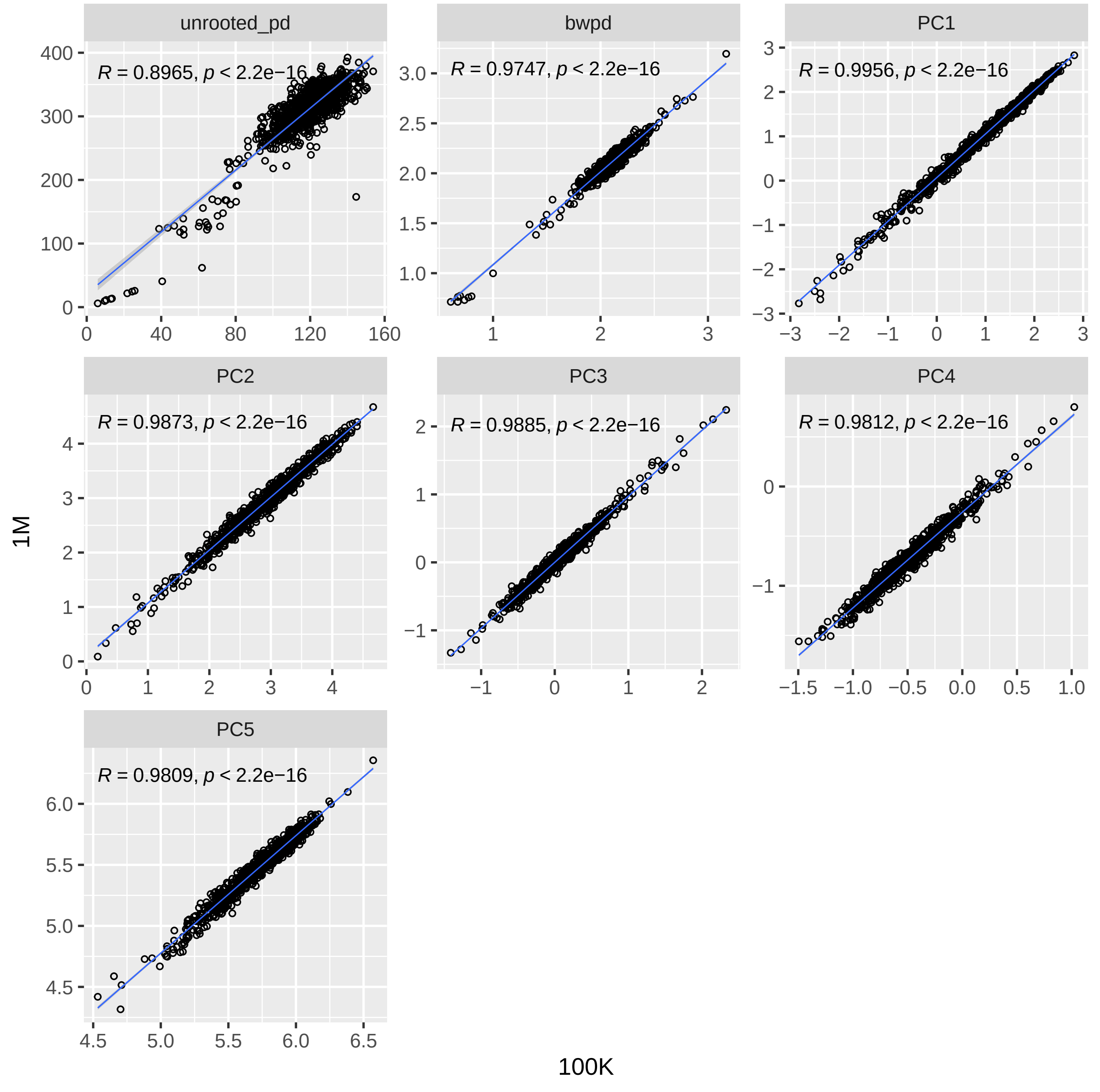
